# Releasing the *octoPus,* an open-source digital tool to promote Integrated Pest Management

**DOI:** 10.1101/2024.08.07.606987

**Authors:** Simone Bregaglio, Eugenio Rossi, Lorenzo Ascari, Gabriele Mongiano, Eleonora Del Cavallo, Sofia Bajocco, Luisa Maria Manici, Antonio Gerardo Pepe, Chiara Bassi, Rocchina Tiso, Fabio Pietrangeli, Giovanna Cattaneo, Camilla Nigro, Marco Secondo Gerardi, Simone Bussotti, Angela Sanchioni, Danilo Tognetti, Mariangela Sandra, Irene De Lillo, Paolo Framarin, Sandra Di Ferdinando, Riccardo Bugiani

## Abstract

Meeting the EU targets to halve chemical pesticide use by 2030 necessitates European farmers to adopt Integrated Pest Management principles as the standard. Decision support systems are valuable tools to meet this target and rely on individual disease models to identify conducive conditions to fungal infections. These models are often proprietary assets of digital startups and agrochemical companies, leading to a lack of transparency for farmers and a bias towards chemical solutions over sustainable practices. We present *octoPus*, the first free digital tool designed to support the control of primary infections of grapevine downy mildew, and we evaluate its performance and behavior on a wide set of environmental conditions and agricultural contexts. We implemented eight models from scientific articles (Rule310, Laore, EPI, IPI, DMcast, UCSC, Misfits, Magarey, the “tentacles”), and evaluated them across Italian grapevine areas from 2001 to 2020. Model outputs were integrated with phenology and susceptibility models (the “eyes”), which were calibrated using data from regional extension services’ bulletins. The simulated infections serve as predictors in a Random Forest algorithm (“brain”) that elaborates an overall risk level (very low to very high). The Llama large language model is used to generate user-supportive messages (the “mouth”).

*octoPus* is released as an open-source software, which reads weather data, executes the models, and presents outputs in natural language and symbolic syntax. Our results showed reasonable accuracy in simulating grapevine phenology (RMSE = 9-10 days) and seasonal risk (RMSE ≈ 0.75). The infection models consistently identified a moisture and thermal north-south suitability gradient in Italy and accurately detected years with low or high downy mildew pressure. However, the models displayed significant differences in the number and dynamics of simulated infections, with two distinct patterns within the ensemble.

By developing and releasing the first free and open-source tool to support the control of grapevine downy mildew, we address a critical gap in the availability and transparency of decision support systems for European farmers. Unlike proprietary models that often lack transparency and may favor agribusiness’ logic, *octoPus* provides a comprehensive and accessible alternative that promotes Integrated Pest Management practices. We propose the adoption of *octoPus* by plant health authorities to identify areas for performance refinement and capabilities expansion.

## 1. Introduction

Supporting the design of evidence-based plant protection strategies requires considering a complex intertwining of biological, economic, and social aspects (Steinberg, 2013). Farmers’ plant protection strategies risk to be dictated by agrochemical companies’ sale efforts rather than being guided by environmental principles (Magarey et al., 2019). Although Integrated Pest Management (IPM) practices have been incorporated into European regulations for years (Directive 2009/128/EC, European Commission, 2009), the proliferation of confusing definitions, inconsistencies in its application, and the limited farmers engagement contribute to the harsh truth that chemical control remains the backbone sustaining plant health programs (Deguine et al., 2021). Additional barriers to IPM adoption are the poor coordination of regional and national programs, the need for technical and educational support to farmers, the translation of plant epidemiology advances into operational services, and, possibly, the cultural backwardness within the agricultural environment (Greitens and Day, 2007; Parsa et al., 2014; Lefebvre et al., 2015).

As a result, it has been demonstrated that pesticide use in the EU increased from 2010 to 2018 due to the member states’ focus on the active ingredients’ risk, rather than pesticide use, and the absence of clear monitoring indicators (European Court of Auditors, 2020; Jacquet et al., 2022). The shortage in funding for public extension services promoting IPM practices and the consequent limited human resources in charge of field scouting worsen the situation. European farmers have therefore increasingly turned to digital startups from private or academic sector to receive technical support for agricultural management. The real concern is that the undisguised objective of these companies is to enter the clutches of multinational agrochemical corporations to capitalize and maximize profit, causing a worrying shift toward market-driven factors in defining plant protection strategies and actual pesticide use (for the US, Wang, 2014; for the EU, Bregaglio et al., 2022).

To promote a wider implementation of IPM practices, the EU Commission has promoted the ’Green Deal’ initiative (2019) and the ’From Farm to Fork’ strategy (2020), aiming at halving pesticide use and associated risks by 2030. Plant disease models are crucial element within these legislations because they can inform farmers on the current weather suitability to pathogen infections, which is crucial to optimize the intervention timing and dosage (Gent et al. 2013; Garcerá et al. 2021). These models are often realized in the form of computer software reproducing the epidemiological cycle of fungal pathogens in response to weather conditions at different levels of complexity, ranging from purely empirical to mechanistic (González-Domínguez et al., 2023). They constitute the main digital asset of public and private Decision Support Systems, where generally a single modelling approach is used to provide infection risk alerts (Chakrabarti and Mittal, 2023). A recent global meta-analysis, drawing from 80 studies spanning diverse pathosystems, proved a significant reduction in pesticide use and disease incidence due to model-based decision support systems compared to standard calendar-based fungicide scheduling (Lázaro et al., 2021). In cases where the disease incidence remained unchanged, the use of models almost halved the median number of fungicide sprays. Their role in meeting EU objectives is even more crucial in organic farming where applications have lower efficacy and need precise scheduling (Tamm et al. 2017).

Current environmental modelling research indicates that multi-model ensembles, i.e., the combined use of alternative algorithms for the same process (Araújo and New, 2007), can outperform the accuracy of single models while allowing the analysis of the uncertainty in predictions (Porter et al., 2014; Sajid et al., 2022). Perspectives and reviews in plant disease also converge on the need of using model ensembles to improve prediction skills (Gessler et al., 2011; Donatelli et al., 2017; Velasquez-Camacho et al., 2023), despite operational realizations still lack. One possible reason is that proprietary plant disease models are considered strategic company assets, so their source code and the protocols used for calibration and validation are rarely released to the community. Consequently, the core equations and the model parameters values often remain undisclosed or incompletely reported in scientific articles, which reduces the likelihood of independent use by third parties. In the meantime, the increasing use of machine learning techniques in plant disease forecast and the recent advancements in Large Language Models (LLMs) potentially open a new era in environmental prediction modelling (Razavi et al., 2022), with hard to predict implications. Combining knowledge from process-based models with machine learning algorithms that can handle large, multi-dimensional datasets has proven to improve accuracy in crop yield forecasting (Bregaglio et al., 2021; Paudel et al., 2022); however, practical applications in plant disease management are still underdeveloped.

In response to these limitations, we developed the *octoPus* (eight – **octo** – models for ***P****lasmopara viticola* **u**nited **s**imulation), a digital tool for the prediction of grapevine downy mildew infections. *octoPus* is composed by a grapevine phenology model to estimate host susceptibility (the “eyes”), a multi-model ensemble to sense the environmental conduciveness to downy mildew infections (the “tentacles”), a machine learning algorithm to synthesize models’ outcomes in the daily level of infection risk (the “brain”) and an open-source LLM which collects information and elaborates support to farmers (the “mouth”). *octoPus* has been calibrated using historical series of phenology and downy mildew infection risk data (Bregaglio et al., 2022b) and has been tested in Italy as the use case to foster its adoption by the farming community *sensu lato*.

## 2. Materials and Methods

The study workflow (Figure 1) has been conceived by ten Italian extension services and the Council for Agricultural Research and Economics which promotes the Misfits initiative, a collective voluntary effort from technicians, researchers, and public officials to build transparent, accessible, and reliable monitoring and modelling tools to promote plant health (Bregaglio et al., 2022a). Step A comprised the implementation of a chilling-forcing model to simulate grapevine phenology, whose outputs have been used in Step B to predict downy mildew susceptibility based on BBCH codes and expert-derived rules (section 2.1). In Step C, eight downy mildew infection models have been reimplemented and homogenised in the outputs format and run in Italy to analyse their behaviour across climatic conditions (section 2.2). The outputs from phenology and disease models have been used as predictors to synthesize the ensemble prediction using a Random Forest algorithm trained on reference infection risk levels (Step D, section 2.3). The information gained analysing models’ behaviour and their daily outcomes have been synthesized in a message processed by the LLM (Step E, section 2.3), to elaborate a tailored decision-support advice for farmers. The *octoPus* source code is freely available at https://github.com/GeoModelLab/octopus.

**Figure 1.**
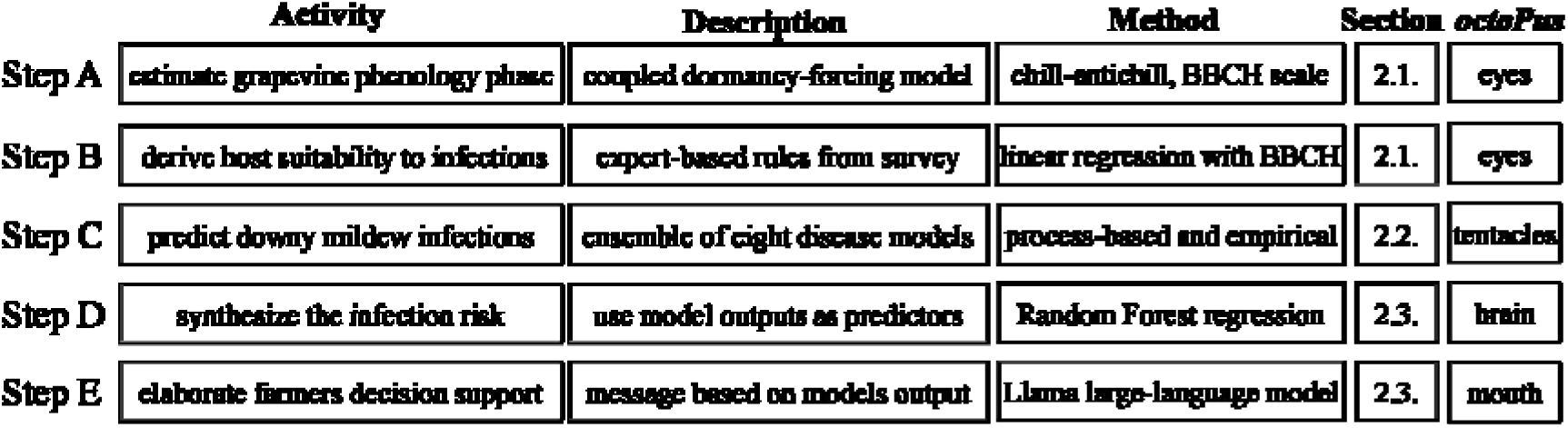
The study workflow divided in five steps, with description of the activity and method, the reference to the paper section, and the corresponding *octoPus* digital components.

Hourly weather data as input for modelling have been downloaded at 0.25° × 0.25° resolution from Copernicus ERA5 dataset (531 grid cells) and referred to the Italian territory in 2000-2021 (Hersback et al., 2023). Weather variables included hourly air temperature (°C), relative humidity (%), precipitation (mm) and leaf wetness (0-1). Hourly vapor pressure deficit (kPa) has been estimated from temperature and relative humidity (Monteith and Unsworth, 1990). Each grid cell has been referred to the corresponding NUTS-2 (region) and NUTS-3 (province) unit using administrative boundaries for subsequent analyses.

### 2.1. Estimating grapevine phenology and host susceptibility

Grapevine phenology has been simulated at a daily time step, coupling a dormancy (Cesaraccio et al., 2004; Didevarasl et al., 2023) and a forcing (Bregaglio et al., 2022a) model. The dormancy model tracks plant progress through endodormancy, considering exposure to low temperatures until the plant’s chilling requirement is met. During ecodormancy, the model uses the degree-day method to accumulate anti-chill days needed to define the bud break date, corresponding to BBCH 11. From that day onwards, grapevine phenology is simulated based on growing degree days accumulation with a non-linear forcing function (Yan and Hunt, 1999) driven by minimum (10 °C), optimum (20 °C) and maximum (35 °C) cardinal temperatures. The grapevine growing season has been encoded according to the BBCH scale from bud break (BBCH 11) to maturity (BBCH 89). The total chilling requirement, as well as the thermal thresholds to trigger BBCH phases have been automatically calibrated using downhill simplex method with a reference dataset comprising 2189 entries from ten Italian regions for the period 2012-2017 (Bregaglio et al., 2022b). Once calibrated on a single ERA5 grid cell at NUTS3 level (province), the phenological model was validated on the other grid cells in the same administrative unit. Host susceptibility to downy mildew infections during the growing season was quantified on a scale ranging from 0% (not susceptible) to 100% (highly susceptible) using BBCH phases as predictors of susceptibility scores based on expert-opinion (Bregaglio et al., 2022a). The information to derive the susceptibility scores from BBCH phases came from surveys to agronomists, officers, and technicians working on extension services with proven experience in disease scouting, plant disease protection, and epidemiology.

### 2.2. Prediction of mildew primary infections

Eight downy mildew primary infection models have been selected from scientific literature (Table 1). Their algorithmic description is provided in Supplementary S1.

**Table 1.**
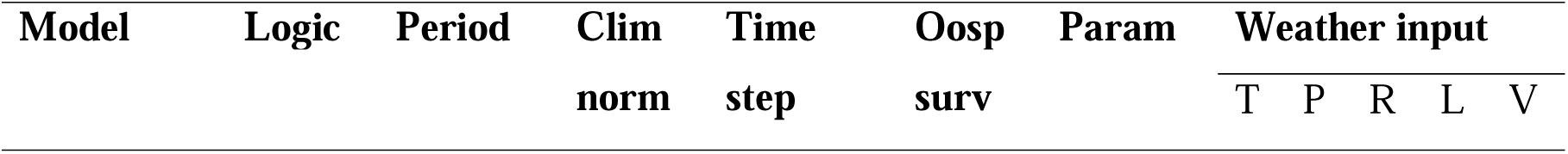
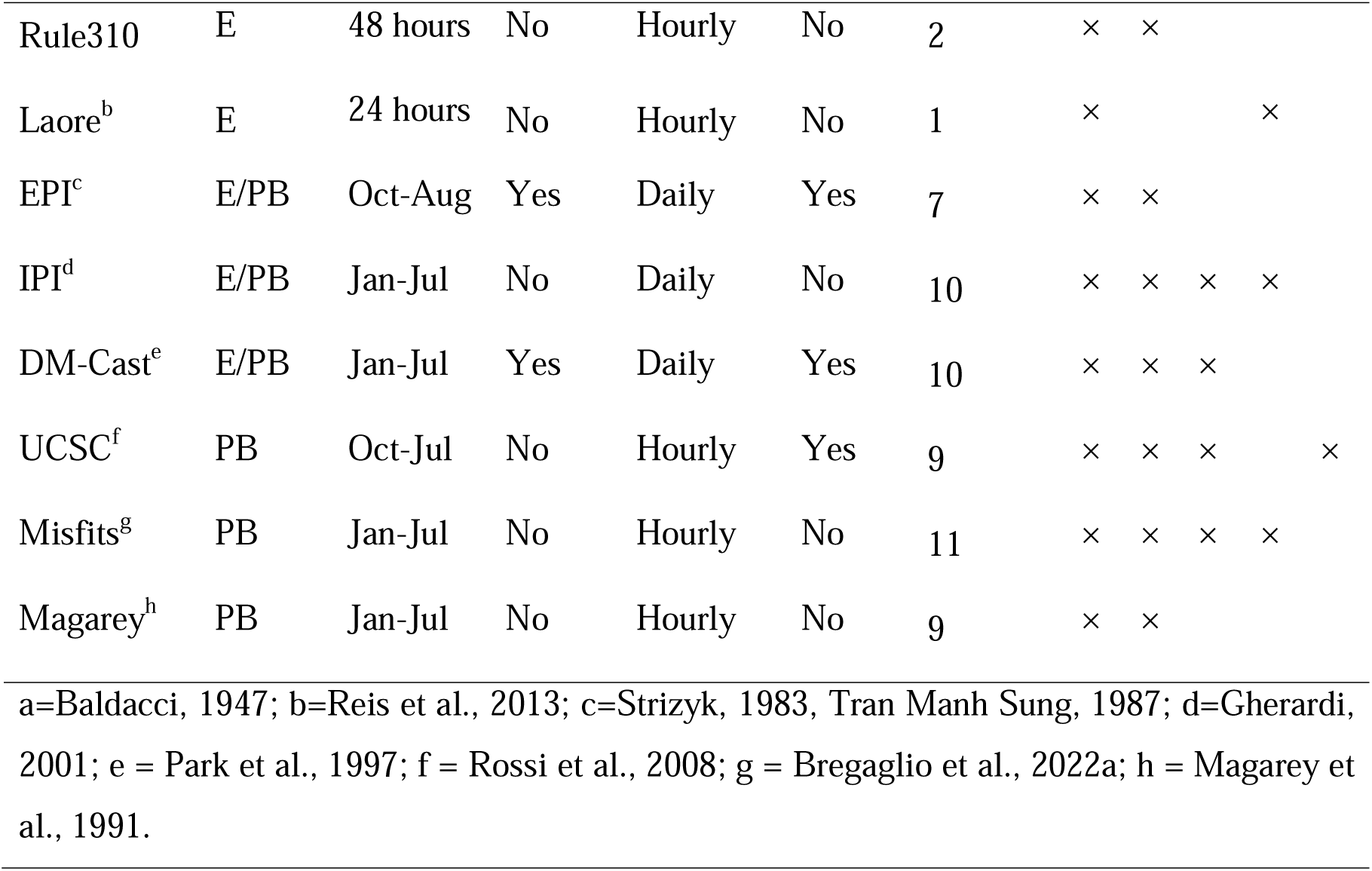
Main characteristics of the eight models for grapevine downy mildew primary infections. E = Empirical; PB = Process Based; months are abbreviated as 3-letter codes; Clim norm = requires climatic norm as input; Oosp surv = accounts for oospores overwintering survival; Param = number of parameters; T = air temperature (°C), P = precipitation (mm), R = relative humidity (%), L = leaf wetness (0-1), V = vapor pressure deficit (kPA). References to scientific articles where the model has been presented are reported below the table.

Two fully empirical models have been implemented, i.e., the Rule310 (Baldacci, 1947), and the Laore model (Reis et al., 2013). The 3-10 ‘rule of thumb’ summarizes a specific set of conditions for the occurrence of primary infections: daily air temperature > 10° C, precipitation > 10 mm in the last 48 hours, and host suitable phenological phase (BBCH > 10). The Laore model triggers a primary infection when an index based on hourly temperature and leaf wetness duration exceeds a user-defined threshold (13 in this study, 12-14 in Reis et al., 2013). Five models do not simulate oospore overwintering, whereas three models explicitly consider this process, i.e., DM-Cast (Park et al., 1997), EPI (Strizyk, 1983; Tran Manh Sung, 1987), and UCSC (Rossi et al., 2008). DM-Cast uses a Gaussian function to represent the probability of oospore germination, whereas EPI considers a potential period (October-March) when oospores are sensitive to weather conditions. These two models work at daily time step and implement functions driven by thermal and pluviometric conditions based on historical series of weather data. UCSC (Rossi et al., 2008) is the more process-oriented model within the ensemble because it reproduces the critical stages of the pathogen’s life cycle at an hourly time step, including oospore dormancy, primary inoculation period, oospore density, germination into sporangia, sporangia survival, zoospore release and survival, infections via zoospores, and the incubation period leading to symptoms onset. The IPI model (Gherardi, 2001) calculates a daily index related to the pathogen infection potential as a function of weather conditions: oospores germination occurs when the cumulated index exceeds a threshold, and primary infections are triggered by simple rules based on temperature, precipitation, and plant phenology. The Misfits model (Bregaglio et al., 2022a) reproduces the weather suitability to sporangia formation, zoospore spread, and consequent primary infections using pathogens’ cardinal temperatures and moisture requirements to modulate functions driven by hourly data. The Magarey model (Magarey et al. 1991) assesses the thermal and pluviometric conditions needed for oospores transport from the soil to the plant, followed by germination and zoospore release when leaf wetness and temperature are conducive, with precipitations leading to a primary infection when leaf wetness duration exceeds 45 degree-hours. The eight models were run considering the period from April 1^st^ to July 31^st^ as representative for the seasonal time window to provide farmers support against primary downy mildew infections, and the daily outputs converted into binary data (0 = no infection day, 1 = infection day; Supplementary S1).

### 2.3. Post-processing models’ outputs and elaborating decision support messages

The host phenological susceptibility score (section 2.1.), the rolling sum of simulated infections in the previous week, and the seasonal total infections (section 2.2) have been used as input to train a Random Forest algorithm (‘randomForest’ and ‘caret’ R packages), using a reference dataset of infection risk. The reference data consisted of 1781 records of downy mildew infection risk on a scale from 1 (very low) to 5 (very high). Six independent evaluators interpreted the information provided in phytosanitary bulletins issued at NUTS-3 (provincial) level and converted them into standardized risk scores (Bregaglio et al., 2022b). The averaged infection risk score from the six evaluators was used as dependent variable to train (70% of the data) the Random Forest regression model using k-fold cross-validation as resampling method (four-fold), after optimizing the ‘mtry’ parameter (6). The remaining 30% of the observations were used for independent validation. The trained Random Forest model (R object) has been imported in C# (R.NET library) and it is executed at runtime using the outputs from the phenology and disease models as predictors. A textual output condensing the study outcomes and models predictions is sent to the Application Programming Interface (API) of the open-source Llama LLM (version llama-2-7b-chat.Q6_K, Touvron et al., 2023). The LLM is used to generate summaries of the models’ outcomes using natural language. The prompt used for this task is reported in Supplementary Materials S2. As a proof of concept, *octoPus* was implemented as a text-based console application. The risk level is displayed using ASCII symbols, colored from blue (very low) to red (very high), and associated with simple 8-bit sounds of increasing frequency according to the infection risk; when conditions are conducive for the infection, the summary text is also generated.

### 2.4. Comparative analysis of models’ behaviour using climatic cluster analyses

Climatic clusters were identified using principal component analysis (PCA) and Hierarchical Clustering on Principal Components (HCPC), utilizing the FactoMineR R package. PCA was conducted using seasonal weather indicators from all ERA5 climate grids, aggregated at the NUTS2 (regional) level. Seasonal weather indicators were derived by averaging seasonal maximum and minimum daily temperatures and relative humidity, and by summing the precipitations and leaf wetness hours. The cumulative number of infections from the eight models was also included as supplementary quantitative variable, which was overlapped to the analysis without contributing to the definition of the principal components space. HCPC was then performed on the first two principal components, with the optimal number of clusters determined according to the hybrid method of Kassambara and Mundt (2020) using the factoextra R package. The daily dynamics of simulated infections was then used to investigate the similarity in models’ behaviour using the Matthews correlation coefficient (φ, Matthews, 1975, eq. 1), which has been computed from binary daily outputs to derive correlation matrices across climatic clusters.

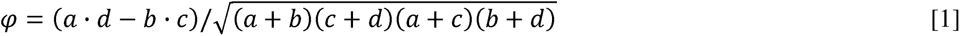

where *a* is the number of 1:1 pairs, *b* is the number of 0:1 pairs, *c* is the number of 1:0 pairs and *d* is the number of 0:0 pairs. A Hierarchical Clustering analysis based on the Matthews Correlation Coefficient has been performed to derive within-clusters dendrograms of models’ similarity based on φ height distance (parallelDist R package). All data analyses were performed using R version 4.4.0 (R Core Team, 2024).

## 3. Results and discussion

### 3.1. Performance in reproducing phenology observations and host susceptibility

The phenology model reproduced grapevine BBCH observations with Root Mean Square Error (RMSE) between 9 and 10 days for leaf development, inflorescence emergence, flowering, and fruit development phases. The maturity phase, which usually occurs in August-September, was characterized by more significant errors (RMSE = 16 days, Supplementary Table S3.1). Despite the timing of BBCH phases exhibited considerable variability across years and NUTS-3 units, our simulations consistently reproduced their trend (Supplementary Figure S3.2). The implementation of the dormancy phase based on the ‘chill-anti chill’ concept (Cesaraccio et al., 2004) improved the prediction accuracy with respect to a precedent modelling work using a thermal time-based model applied on the same dataset (Bregaglio et al., 2022a) in Veneto (RMSE 8.3 vs. 10 days), Emilia-Romagna (RMSE = 8.7 vs. 10.5 days), Sardinia (RMSE 9.4 vs. 10.2 days) and Basilicata (RMSE = 10 vs. 12.7 days). These errors align with phenology modelling studies conducted in Mediterranean conditions (Ortega-Farias and Riveros-Burgas, 2019) and other grapevine-growing environments (Chile, Garcia-Gutiérrez and Meza, 2023; China, Wang et al., 2020). These studies focused on widely grown grapevine varieties in Italy (Cabernet-Sauvignon, Merlot, Chardonnay) and addressed model performances with variety-specific parameter sets. Here, we used a single parameter set calibrated at the NUTS-3 level; users can customize phenology parameterizations to tailor *octoPus* application for different varieties and environments. The linear interpolation of the scores of host susceptibility to downy mildew derived from expert evaluators’ interpretation of phytosanitary bulletins (Bregaglio et al. 2022a, see Methods section 2.3) determined a gradual increase from BBCH 11 to 15 (first leaf unfolding to shoots reaching 10 cm), followed by a sharp rise towards BBCH 61 around mid-May (start of flowering). The host susceptibility markedly declined in all study areas after mid-June, along with the progression of fruit development stages. The proposed method to derive host susceptibility based on host phenology gives a simple but robust representation of grapevine susceptibility, which aligns with common knowledge (Bugiani and Bariselli, 2022) and epidemiological studies, e.g., the evidence of host-pathogen phenology synchrony mechanisms during oospore maturation and germination (Maddalena et al., 2021).

### 3.2. Similarities in primary infections dynamics across climatic conditions

The results of the multivariate analyses on seasonal weather indicators are shown in Figure 2.

**Figure 2.**
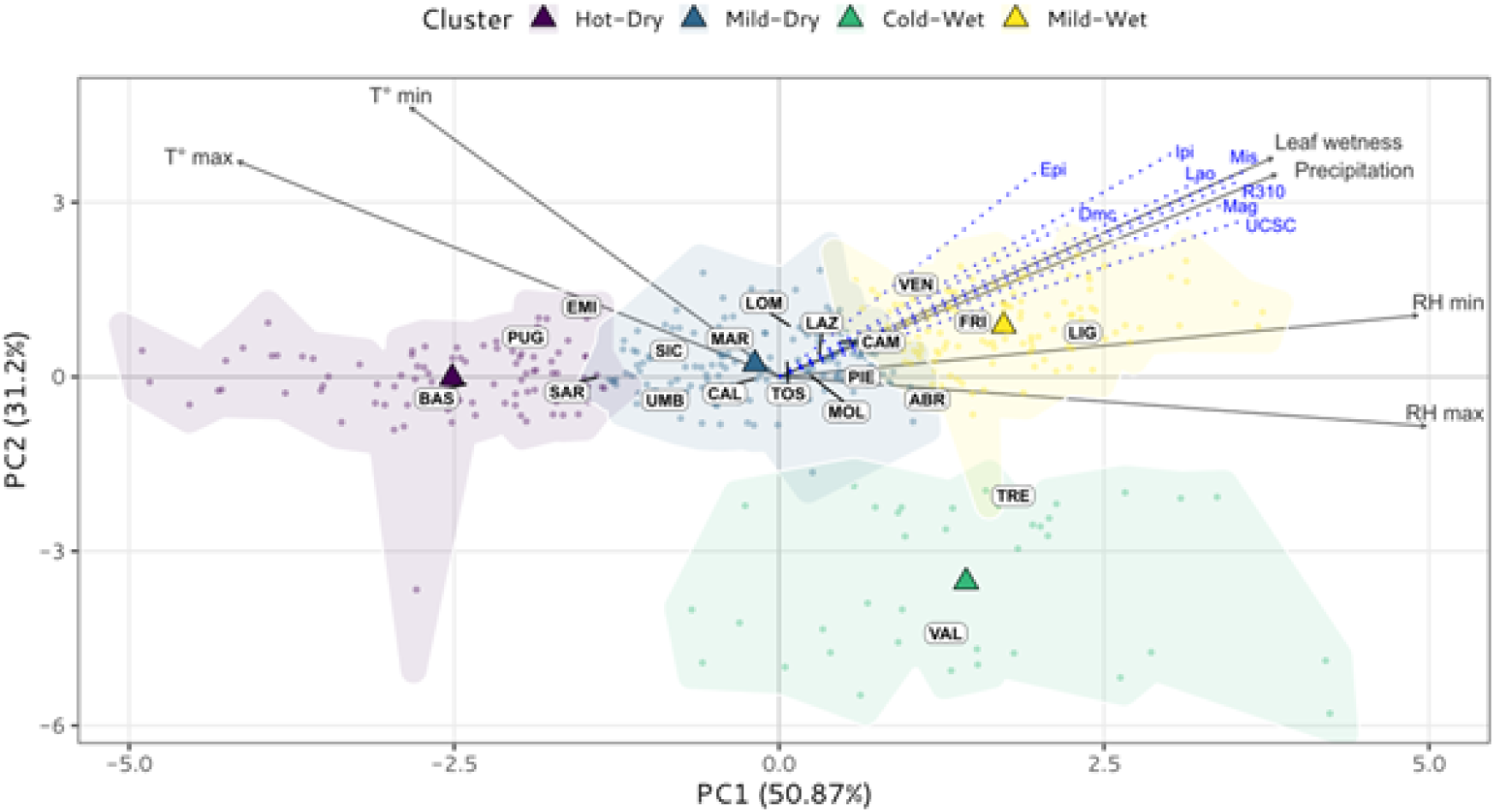
Results of the Principal Components (PC) and Hierarchical Clustering Analyses conducted on the seasonal averages (May-July) of maximum (T° max) and minimum (T° min) air temperature (°C), maximum (RH max) and minimum (RH min) relative humidity (%), cumulative leaf wetness (hours), and precipitation (mm) grouped by region (NUTS-2). Vector endpoints indicate the strength and direction of correlation between the variables and the PCs. Each point represents a season from 2000 to 2021, color-coded by cluster membership, with cluster centroids indicated by triangular shapes.

The supplementary variables representing simulated seasonal infection predictions from the eight infection models (Table 2) are projected onto the PC space (in blue). The first two Principal Components (PCs) explained 82.1% of the original data variability and were considered in the analysis (Supplementary S4). The Hierarchical Clustering on Principal Components (HCPC) analysis led to four clusters, which have been encoded as Hot-Dry, Mild-Dry, Cold-Wet and Mild-Wet, based on their temperature pattern (Hot, Mild, or Cold) and precipitation regime (Wet or Dry). The mountainous regions of Valle d’Aosta (VAL) and Trentino-Alto Adige (TRE) in northern Italy are in the Cold-Wet cluster, which is associated to low temperature (8 °C) and precipitations (172 mm) in April-July. The other NUTS-2 units are distributed along PC1 from negative (Hot-Dry) to positive (Mild-Wet) coordinates and follow a clear south-north gradient, corresponding to increasing precipitation (Hot-Dry = 158 mm, Mild-Wet = 385 mm) and Leaf Wetness Hours (Hot-Dry = 511, Mild-Wet = 987), but decreasing seasonal maximum temperatures (Hot-Dry = 24 °C, Mild-Wet = 22 °C). The supplementary variables representing simulated seasonal infection predictions from the eight models are positively associated to both PC1 and PC2 (Figure 2). These results suggest that *octoPus* models tendentially simulate the largest pathogen pressure in correspondence to mild temperatures and elevated moisture, converging in their predictions. Major limiting factors to pathogen infections were cold temperatures in mountainous areas (Cold-Wet cluster) and low moisture and precipitations (Hot-Dry) in the southern regions. Simulated seasonal dynamics of downy mildew infections are shown in Figure 3, grouped by the four clusters, along with the host susceptibility score computed from phenology (section 3.1). The comparison of daily infection dynamics reveals diverging behaviors among the models, even though they exhibit similar relative sensitivity to the climatic clusters. The number of seasonal infections depicted the following descending order in the models: DM-Cast > IPI > EPI > UCSC > Rule310 > Laore > Misfits > Magarey. In the Mild-Wet cluster, some models (DM-Cast, IPI, EPI, UCSC, Laore) agreed in simulating a very low weather conduciveness to infections until mid-May, followed by a steep increase of the pathogen suitability onward, whereas Rule310, Misfits and Magarey steadily predicted an earlier onset of conducive conditions. The average seasonal infections in this cluster ranged between 7.6 ± 2.5 for Magarey to 24.6 ± 5.5 for EPI. All models consistently simulated an earlier onset of downy mildew infections in the Mild-Dry and the Hot-Dry clusters, because of warmer temperatures. In these clusters, the rate of simulated infections decreased after June in relation to lower precipitations and drier conditions.

**Figure 3.**
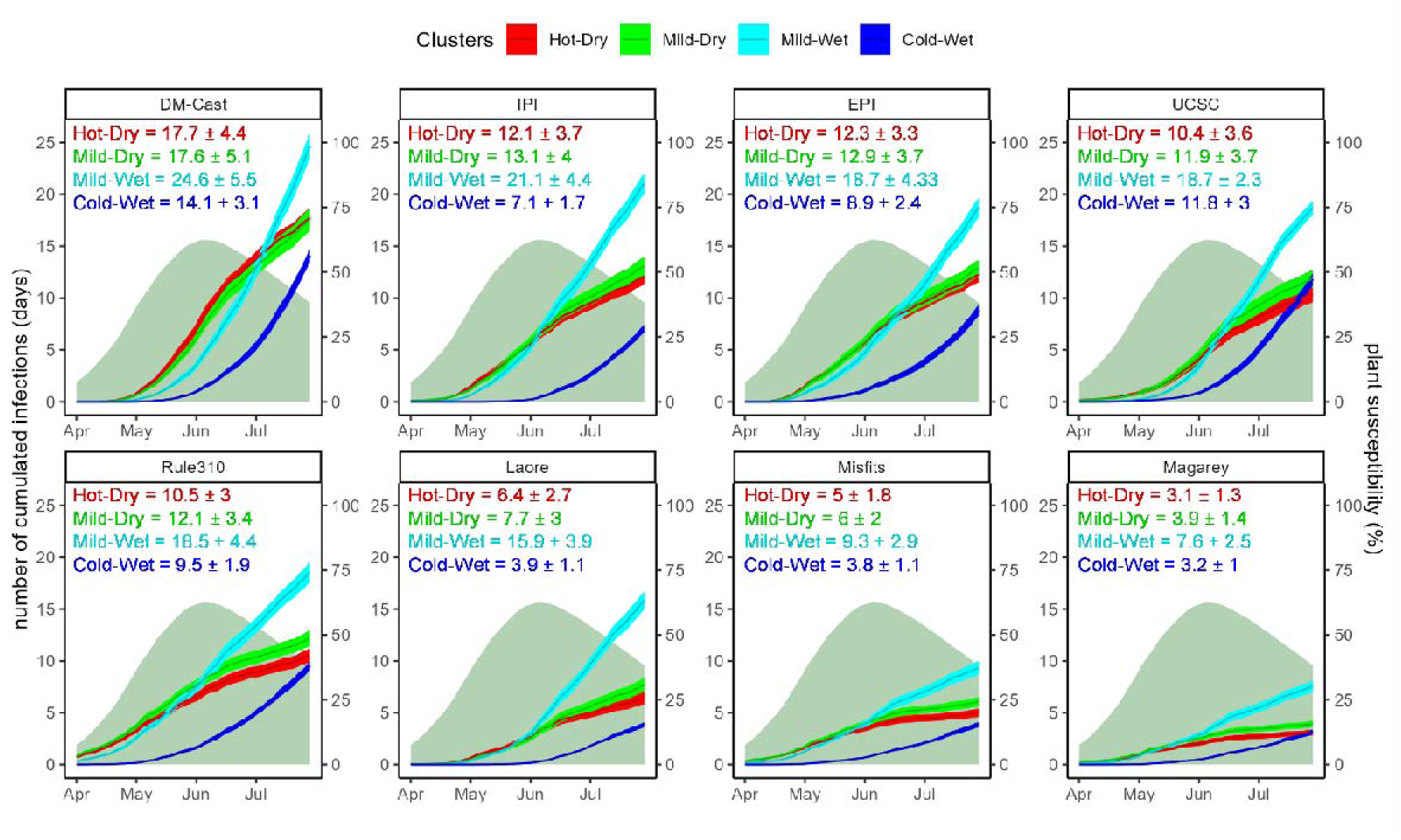
Cumulated seasonal infections simulated by the eight models (colored lines and shades, primary y-axis) and host susceptibility scores (green area, secondary y-axis) in the four climatic clusters. The line is the average cumulated number, and the shades correspond to the mean ± one standard error. The text in each facet reports the average seasonal infections ± one standard deviation.

All models simulated a similar number of infections in these clusters, with a maximum difference of two days for each model. The models behavior also converged in the Cold-Wet cluster, where infections were markedly delayed and started to be simulated from mid-May. However, the absolute number strongly varied from less than 5 (Laore, Misfits, Magarey) to more than 10 (DM-Cast, 14.1 ± 3.1; UCSC, 11.8 ± 3). The Hierarchical Clustering conducted considering φ distances in the four climatic clusters depicts two main behavioral patterns among models (Figure 4). The first group of similarity comprised Misfits, Rule 310 and Magarey, which consistently showed lower internal distance (φ < 0.6), and higher differences (φ > 0.8) with the other five models. Within this group, Magarey mostly differed in the Mild-Dry and Hot-Dry clusters, whereas Misfits deviated to the other two models in the Cold-Wet and Mild-Wet areas.

**Figure 4.**
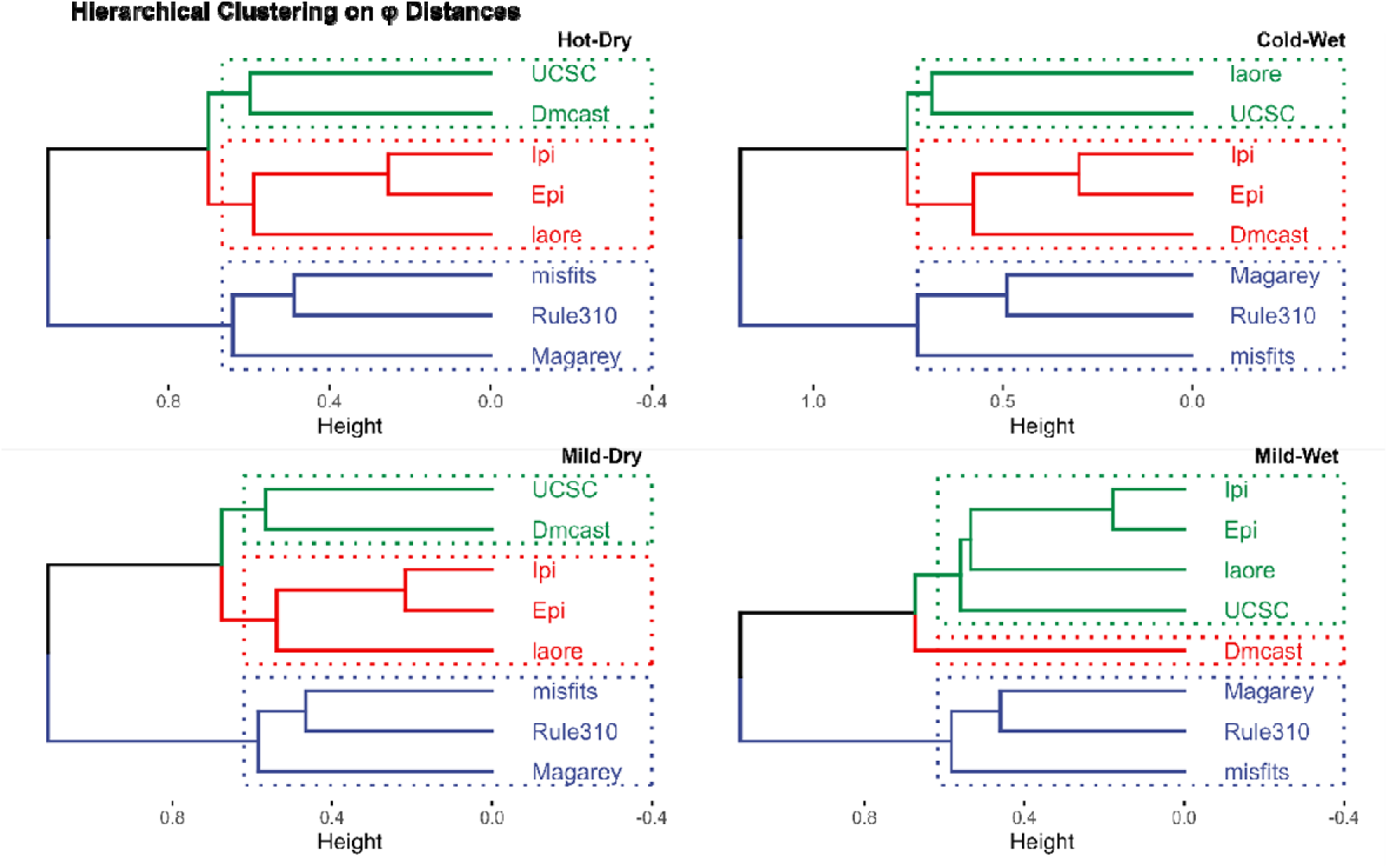
Dendrograms of models’ similarity based on the correlation matrices derived from daily simulated series of binary data (0 = no infection, 1 = infection). Four dendrograms are reported, one for each climate cluster from the multivariate climatic analyses.

We decided to further highlight models’ differences in the second group, which included all the other models. IPI and EPI showed higher similarity (φ < 0.4), followed by Laore. UCSC and DMcast also showed an overall similar behavior (φ < 0.7). The large differences in the absolute number of infections simulated by the eight models (Figure 4) highlights the appropriateness of opting for an ensemble where the single models’ performance is investigated to better interpret the outputs and translate them into decision support. This also reinforces the need for sharing experimental field datasets where single models can be jointly calibrated to refine their performances, which is a recognized limitation in studying plant disease epidemics (Savary et al., 2019). A comparison study of four downy mildew infection models (Rule310, EPI, Dm-Cast, UCSC) has been performed by Caffi et al. (2007), who applied them on a single location in Sardinia (Hot-Dry cluster) from 1996 to 2004. They found that UCSC outperformed the other three models, and suggest to discard Rule 310 and EPI as too simplistic in their formulation. In our study, the seasonal infections simulated by these four models were comparable (Figure 3), whereas Magarey and Misfits tended to detect lower infections. With the tested parameterizations, correlation analysis showed similarities between the UCSC and DmCast models (Figure 4), whereas the two more ‘conservative’ Magarey and Misfits aligned with Rule-310, creating two distinct behavioral patterns.

Developing multi-model ensembles where alternative approaches are run in parallel has become a standard practice in environmental modelling (e.g., crop yield forecasting, Martre et al., 2015; biogeochemical science, Sandor et al., 2018) but it is still widely unexplored in plant disease forecasting. The primary aim of decision support systems targeting plant diseases should be lowering down the number of treatments during the season (Gherardi 2001). This is especially valid in the case of *Plasmopara viticola*, where fungicide spraying occurs every 7-10 days from April to August, leading to 12-16 fungicide treatments per year (Cabús et al., 2017). One of the main concern in ensemble modelling is that including very similar models risks to place undue weight on a single approach (Wallach et al. 2014). However, the relationship between algorithmic formalization and simulated outcomes is not straightforward, given that the same model with different parameterization can lead to very different results (Li et al. 2015; Folberth et al. 2016). In our study, EPI and IPI are both negative prognosis models that produce very similar results although they employ different calculations to determine the critical threshold triggering infections. For practical reasons we run the models with default parameters, leaving the specific parameterization to users and developers for *octoPus* application in operational contexts. Performing recursive calibration of the parameters of each infection model is indeed possible and may be needed when used in real-world applications. The parameters are stored in a .csv file with minimum, maximum, and unit of measurement, and their values are configurable by the user. Relying on ‘sensitive’ models which tend to simulate a large number of infections could lessen the benefit in fungicide sprays savings, with respect to simple calendar-based control systems. On the other hand, adopting ‘conservative’ models, which underestimate the infection risk, is likely to result in insufficient protection, potentially leading to poor control and higher yield losses. Models biased into predicting more infections could be used as sensitive alerts during high risk periods, while environment-specific knowledge and users experience could allow changing the models weights to customize the plant protection strategy.

### 3.3. Spatial and temporal variability of pathogen pressure across Italy

The average seasonal downy mildew infections simulated in Italy from 2001 to 2020 followed a North-South latitudinal gradient (Figure 5). The eastern and northern regions experienced more than 15 infection days per season on average, while southern areas and islands have been associated with less than 10 seasonal infections in most years, consistently with empirical evidence (Galassi, 2023). In the northern regions near the Alpine arc, a large number of primary infections were simulated in warmer seasons, suggesting that in these areas low temperature may be the main inhibiting factor for the pathogen. A very high pathogen pressure has been simulated in 2002, 2014, 2016 and 2018. In these years, the spatial heterogeneity decreased, with an average of more than 14 infection days in 8 regions out of 20, and with only Valle d’Aosta (VAL, Cold-Wet cluster), Sicily and Calabria (SIC, CAL, Hot-Dry cluster) associated to less than 10 infection days (Figure 5). Focusing on Northern Italy, the longitudinal gradient varied across years, with western regions characterized by higher infections in 2008 and 2018, and eastern regions more affected in 2013, 2014, 2016 and 2020.

**Figure 5.**
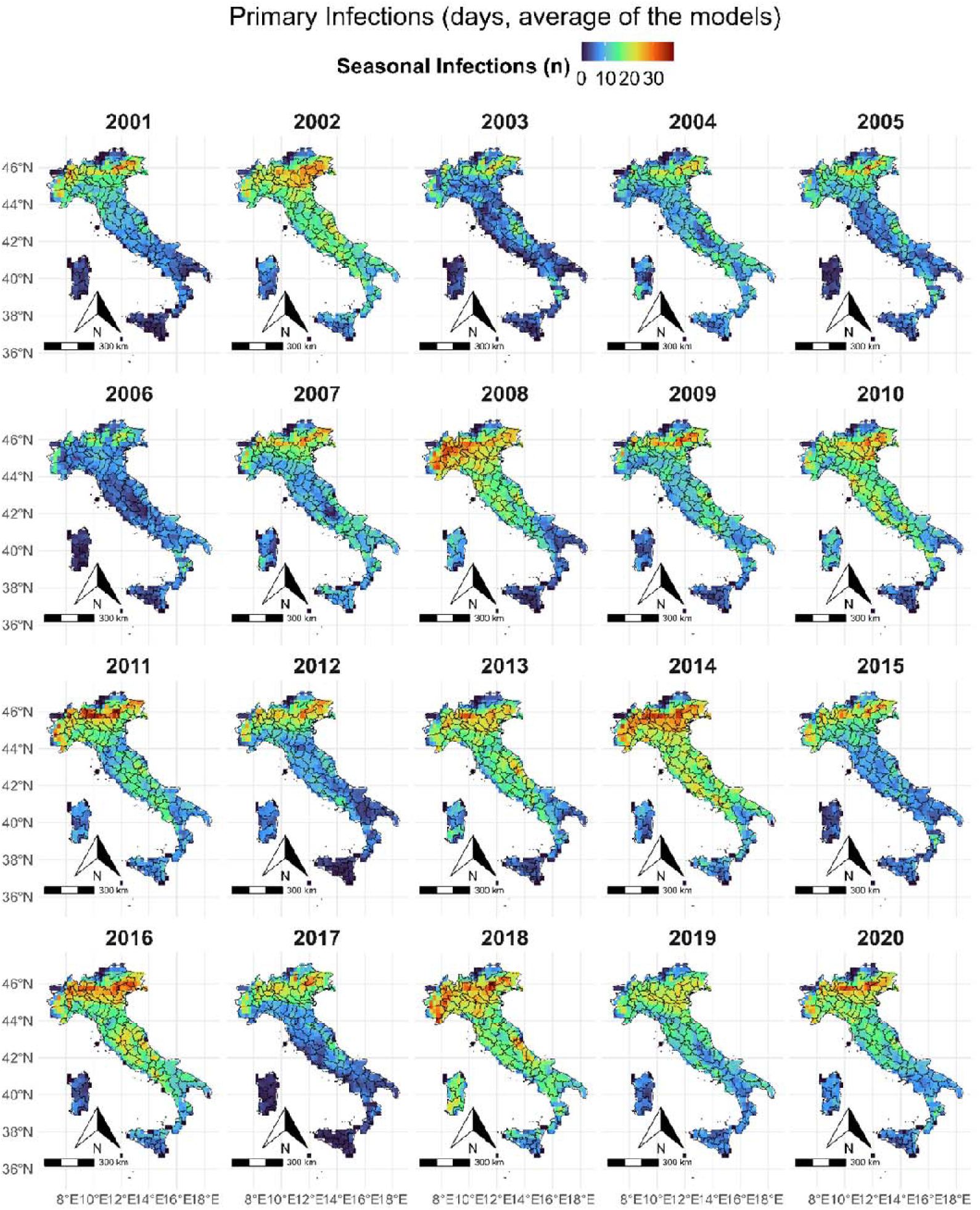
Average number of seasonal simulated downy mildew infections from the eight models in the period 2001-2021 according to the ERA5 grids (0.25° × 0.25°).

The *octoPus* models agreed in simulating very low pathogen pressure in 2003, 2005, 2006, and 2017, with less than 9 seasonal infections in the whole area. The spatial heterogeneity increased in low conducive years, where seasonal infection hotspots were mostly placed in northern and coastal eastern regions. During these years, Marche and Abruzzo (Mild-Dry cluster) showed similar infections than Veneto and Friuli-Venezia Giulia (Mild-Wet cluster). The ensemble’s simulated infections mean agrees with prior modelling and experimental studies, where the total downy mildew primary infections ranged from 3 to 8 in Emilia-Romagna (Mild-Dry cluster, 1995–2006, Rossi et al., 2008), and from 3 to 29 in Tuscany (Mild-Dry cluster, 1995–2003, Dalla Marta et al., 2005). Despite official data on the actual historical pathogen pressure being not available, national web magazines and conference papers agree in indicating 2014 (Mazio et al., 2014), 2016 and 2018 (D’Ascenzo and Crivelli, 2023; FitoGest, 2024; Moser, 2024) as very conducive years for downy mildew primary infections, and 2003 and 2017 as seasons characterized by exceptionally low pathogen pressure (Borgo, 2014; Galassi, 2018).

### 3.4. Elaboration of decision support based on models’ outputs

The Random Forest (RF) model trained using the ensemble outputs as predictors of the seasonal infection risks from phytosanitary services led to balanced accuracy in cross-validation (RMSE = 0.77, mean absolute error, MAE = 0.63 and R^2^ = 0.42). When applied on the independent validation dataset (30% of the data), the performance was comparable (RMSE = 0.74, MAE = 0.61, R^2^ = 0.46). The relative importance of the predictors highlighted host plant susceptibility as the most relevant variable (100), followed by the seasonal infections simulated by Rule-310 (82.7), Misfits (71.2), and Magarey (41.1) models. Considering the rolling sum of infections, DmCast (22.1) and Ipi (17.9) were the most relevant contributors (Supplementary Table S5). Supplementary Figure S5.1 presents the outputs of the Random Forest in comparison to the reference risk level and the simulated infection dynamics. The machine learning model captured the variability of the reference infection risk with significant differences between years and regions. The comparison of the RF model outputs with the observed risk level denotes a correct reproduction and highlights significant differences between years and regions. The RF model accurately identified 2013 and 2016 as the most conducive years for infections and 2015 and 2017 as the ones with lowest pathogen pressure. The simulated risk also captured macro-differences between Italian regions, with Sardinia (Hot-Dry cluster) and Veneto (Mild-Wet cluster) being at the extremes.

The study outcomes have been used to generate natural language summaries using a Large Language Model (Supplementary Material S2), prompted with the outputs from all *octoPus* models (grapevine phenology, host susceptibility, primary infections, and Random Forest).

The *octoPus* console application reads weather data (sample files are provided for each Misfits region), executes the workflow (Figure 1) and delivers synthetic information in the form of natural language, symbols, and sounds (Figure 6, Supplementary Video S6). Users can set the thresholds correponding to the risk level from the Random Forest and to the number of concordant models simulating a primary infections to trigger the call to the LLM.

**Figure 6.**
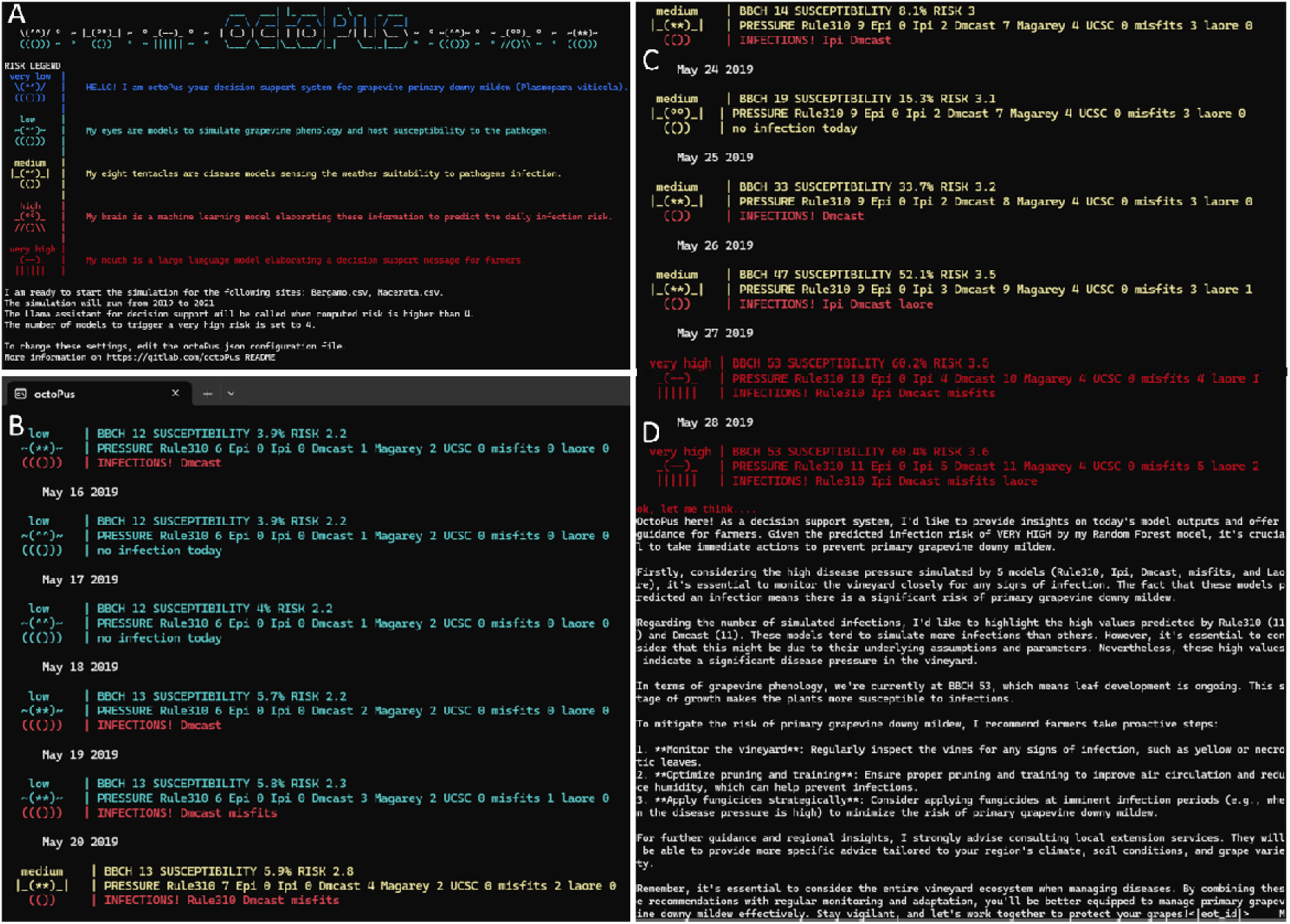
The *octoPus* console application opening screen (A) with symbol and color legends, and simulation settings (sites, years, Llama assistant parameters); simulation outputs in Bergamo, 2019 from May 15 to 20 (B) and May 23 to 27 (C); the message written by the Llama virtual assistant on May 28 (D).

The *octoPus* simulation in Bergamo, 2019 (Lombardy region, Mild-Wet cluster) outlined a low downy mildew infection risk from May 15 to 19 (BBCH 12-13), with only DM-cast signaling three infection events on May 15, 18 and 19, when also Misfits agreed (Figure 6B). Then, from May 20 to 26 the predicted risk increased to medium, and also Laore, Rule310 and IPI started predicting infections (Figure 6C). The grapevine phenology reached BBCH53 (visible inflorescence) on May 27, when host susceptibility increased and five models predicted a downy mildew infection (Figure 6C). These conditions trigger the generation of a decision support message, which is displayed to the user (Figure 6D): the generated message outlined the ongoing scenario on grapevine phenology and pathogen pressure, and discussed the models outputs in relation to their similarity and behavior. Finally, it advised the user on proactive management to control downy mildew, suggesting to consult regional extension services to receive advanced technical support (Figure 6D). The capabilities of LLMs are expanding rapidly, especially in medical sciences, where clinical decision-making and psychological care can be enhanced and fastened (Teixera-Marques et al., 2024), although traditional expert analysis in complex clinical scenarios is still required (Sandmann et al., 2024). The potential of LLMs to improve decision support systems in plant pathology and in other agricultural domains as crop yield forecasting, irrigation planning, precision farming and supply chain optimization is still untapped (Gaddikeri et al., 2023), despite pioneering applications have been proposed (Qing et al., 2023). A limitation is that virtual assistants encounter skepticism and resistance, primarily due to concerns over data privacy, reliability, and model biases. Here we choose to rely on Llama, a research-driven and open-source LLMs rather than other propietary solutions, in line with the willingness to promote *octoPus* as free software. By analyzing models outputs based on the outcomes of this study, the LLM was capable of providing an overview of pathogen pressure and grapevine phenology, offering farmers basic information to implement management strategies. An expert-based LLM training on the best plant protection strategies based on real time conditions could improve the quality of the support, as well as the implementation of chat-like capabilities to enhance users experience.

## 5. Conclusion

This paper releases to the community *octoPus*, a digital tool to assist its users in grapevine downy mildew primary infections control. The design of the conceptual workflow has been participatively realized with officers and technicians from extension services, who set the priorities. Together, we created the *octoPus* to address these objectives through implementation in an open-source software, and we released it to foster collaborative improvements from third parties. Our findings are meant to benefit technicians in drafting guidelines and bulletins, as well as farmers who want to apply plant protection products precisely where and when needed, aligning with IPM principles and regulatory requirements. Future research will need to further verify these results with field experiments to refine the models calibration and the ensemble approach, aiming at giving the octoPus trust, which can be built only with reliable predictions.

## Acknowledgments

The Authors gratefully thank all the people participating to the MISFITS initiative for their active contribution to the discussions which shaped the modelling workflow presented here, and for their precious feedbacks on the priorities and needs for further improvements.

## Authors’ contribution

All authors contributed to the study conception and design. The code implementation was performed by Simone Bregaglio and Lorenzo Ascari. Data analyses were performed by Simone Bregaglio, Lorenzo Ascari and Gabriele Mongiano. The curation of the git repository has been carried out by Eugenio Rossi, Eleonora Del Cavallo and Gabriele Mongiano. The first draft of the manuscript was written by Simone Bregaglio. Eugenio Rossi, Eleonora Del Cavallo and Gabriele Mongiano commented on previous versions of the manuscript. All authors read and approved the fnal manuscript.

## Code Availability

The octoPus source code is freely downloadable at https://github.com/GeoModelLab/octopus

