## Supplementary S1 for "Releasing the *octoPus,* an open-source digital tool to promote Integrated Pest Management"

Supplementary S1. Algorithmic description of grapevine downy mildew primary infection models

Eight models designed to predict primary infections of grapevine downy mildew (*Plasmopara viticola*) were selected from the scientific literature.

- **Rule of three-tens** (*“Regola dei tre dieci*”, Baldacci 1947). This is a simple empirical model that identifies the three main drivers for the initiation of primary *P. viticola* infections: daily air temperature > 10°C, precipitation of the last 48 hours > 10 mm, and a suitable host phenological phase BBCH > 10.
- **DMCAST** (Downy Mildew foreCAST model, Park et al. 1997). This is a model able to predict both primary and secondary *P. viticola* infections but, in the scope of the *octoPus,* only the equations governing primary infections were considered. One of the model’s assumptions is that oospores maturation follows a normal distribution described by:

$$P\left( d \right)=\frac{1}{\sigma\cdot\sqrt{2\pi}}\cdot e^{\frac{{-\left( d-\mu\right)}^{2}}{{2\sigma}^{2}}}$$

Where $P\left( d \right)$ is the probability of oospores to reach maturity at day $d$, where $d$ > January 1^st^. The parameters $\mu$ and $\sigma$ are the average and the standard deviation of the number of days necessary to reach oospores maturation and are calculated as:

$$\mu=118-0.3\cdot Ra$$

$$\sigma=13.5+0.02\cdot Ra$$

$Ra$ represents the effect of the cumulative daily precipitation sum calculated at January 31^st^ with the equation of Tran Manh Sung (1990). This approach postulates that daily precipitation between September 21^st^ and January 31^st^ has a direct effect on oospores maturation. The effect can be categorized as positive (POS) or negative, caused either by lack (LAC) or excess (EXC) of precipitation:

$$Ra=\sum_{21set}^{31gen} POS-\left| EXC-LAC \right|$$

To calculate this effect, the daily precipitation, $Rd$ (mm), is used, together with: the average daily precipitation over a period of 30 year ($RM$, in mm), a minimum threshold below with oospores’ maturation undergoes a negative trend (following the relation: $Hm={RM}/{RDM}$, where $RDM$ is the 30-years average of monthly rainy days), and a maximum precipitation threshold, $HM$ above which a negative effect on the maturation of oospores is applied through the following equation: $HM=RM+{sdRM}/{RDM}$.

Given that, for every day:

$$se Hm<Rd\leq HM\underset{\to}{}POS\left( d \right)=Rd$$

$$se Rd<Hm\underset{\to}{}LAC\left( d \right)=Hm-Rd$$

$$se Rd>HM\underset{\to}{}EXC\left( d \right)=Rd-HM e POS\left( d \right)=HM$$

Through these calculations, the model returns a ‘threshold of attention’ corresponding to the day at which the cumulative proportion of mature oospores reaches 3%, after which they are ready to germinate. Oospores sporangia germination is triggered every time T > 11°C and precipitation > 2 mm. At T > 11°C, seven to twelve days are needed to complete sporangia germination cycle, after which they survive for ≈10 days. For every germinated sporangium an infection process starts every time T > 11°C and precipitation > 0.2 mm.

- **EPI** (*Ètat Potentiel d’Infection*, Strizyk (1983) and following upgrades, see Tran Manh Sung (1987)). The model runs from October 1^st^, date when oospores formation begins, to August 31^st^, after which infection risk is negligible due to harvest. In the model timeframe, two distinct periods are considered: a winter period, corresponding to a potential energy phase of the pathogen, and a vegetative period, corresponding to pathogen’s kinetic energy phase.

The potential energy phase ($Pe$) goes from October 1^st^, day 1 of the model simulation, to March 31^st^, day at which the capacity of mature oospores to produce aggressive zoospores is estimated. This ‘fecundity’ is calculated every ten days starting from a value set to 0 at the end of September, following the equation:

$$Pe=\left[ 2\cdot ct\cdot\left( \sqrt{H}-\sqrt{Hc} \right) \right]$$

$$+\left[ 0.2\cdot ct\cdot\left( \sqrt{H}\cdot\sqrt{T}-\sqrt{Hc}\cdot\sqrt{Tc} \right) \right]$$

$$-\left[ \left( {NGm\cdot1.5}/{18} \right)\cdot log\left( {Hd}/{NGd} \right) \right]$$

Where, $Hm$ is average monthly precipitation over a period of 30 years (mm), $H$ is the monthly precipitation (mm), $Hd$ represents the precipitations of the decade (mm), $Tm$ is the average monthly temperature over a 30 years period (°C), $T$ is the average monthly temperature (°C), $NGm$ is the monthly number of rainy days over a period of 30 years, $NGd$ is the number of rainy days in a decade, $Hc=Hm \times0.95$ representing the monthly critical rainfall (mm) and $Tc=Tm\times0.95$ representing the critical monthly temperature (°C), and $ct$, which is a monthly ranking coefficient that is set to 0.2 for October and November, 1 for December, and 0.8 for January, February, and March.

The kinetic energy phase $\left( Ke \right)$ starts on April 1^st^ until the end of August. It describes the infective period of the fungal pathogen and is described by the daily infection capacity, calculated as:

$$Ke=0.012\cdot\left[ \frac{\left( \frac{5\cdot RHMn+3\cdot RHg}{8} \right)^{2}\cdot\sqrt{Ti}-{RHg}^{2}\cdot\sqrt{Tm}}{100} \right]$$

Where $RHMn$ is the average night-time relative humidity over a period of 30 years (%), $RHg$ is the average daily day-time relative humidity (%), $Tm$ is the average monthly temperature over a period of 30 years (°C), and $Ti$ is the average daily temperature (°C).

From $Pe$ and $Ke$ the $EPI$ (*Ètat Potentiel d’Infection,* potential infection state) is derived in the following way:

$$EPI=\sum_{October}^{March} Pe+\sum_{April}^{Spetember} Ke$$

New infections are triggered when $EPI>0$ and simple eco-physiological thresholds, like temperature, precipitation, and plant phenology, are met i.e., using the rule of three-tens.

- **IPI *Plasmopara viticola*** (*Indice di Potenziale Infettivo,* Index of Potential Infection). Is a negative prognosis model developed by the phytosanitary service of Emilia Romagna region, Italy (Gherardi 2001). The model calculates a daily potential infection index. When its cumulative sum exceeds a pre-alert threshold, primary infections of *P. viticola* can occur. Primary infections are calculated following simple eco-physiological rules, i.e., the rule of three-tens. Additional indices are required to calculate the IPI:
  - Average temperature index $(IndTmed)$, which can assume only values between 0 and 1 $(IndTmed<0\underset{\to}{}0, IndTmed>1\underset{\to}{}1)$:

$$IndTmed=a\times(-2.19247+0.259906\times Tm-0.00013\times{Tm}^{3}-6.095832\times{10}^{-6}\times{Tm}^{4})$$

Where $Tm$ is the average daily temperature (°C), $Tmin$ is the minimum daily temperature (°C), and $a$ is a correction index valorised according to the following conditions:

$$if Tmin>13^{\circ}C\underset{\to}{}a=1$$

$$if Tmin\geq13^{\circ}C\underset{\to}{}a=0.35+0.05\cdot Tmin$$

- $(IndPrp)$, Precipitation index, which can only assume values between 0 and 3 $(IndPrp<0\underset{\to}{}0, IndPrp>3\underset{\to}{}3)$:

$$IndPrp=\left( 0.00667+0.194405*{Prp}_{48}+0.0002239*{{Prp}_{48}}^{2} \right)$$

Where ${Prp}_{48}$ is the cumulative precipitation sum of the last 48 hours (mm).

- Leaf wetness index $(IndBf)$, that can only assume values between 0 and 1 $(IndBf<0\underset{\to}{}0, IndBf>1\underset{\to}{}1)$, and whose value is calculated only when daily precipitation is greater than 0.2 mm and leaf wetness is present for more than 5 consecutive hours:

$$IndBf=0.004*{Bf}^{2}+0.008*Bf-0.01$$

Where $Bf$ are the hours of leaf wetness in a day (h).

- Relative humidity index $(IndUr)$, whose values are only between 0 and 1 $(IndUr<0\underset{\to}{}0, IndUr>2\underset{\to}{}2)$:

$$IndUr=(-69.994545+1.502*Urmed-0.007818*{Urmed}^{2})/2$$

Where $Urmed$, is the daily relative humidity (%).

The $IPI$ index is finally calculated in the following way:

$$if {Prp}_{24}\neq0\underset{\to}{} IndTmed\cdot(IndBf+ IndPrp)$$

$$if Tmin>9^{\circ}C\underset{\to}{} Se IndTmed\cdot\left( IndBf+ IndPrp \right)>IndTmed\cdot IndUr\underset{\to}{}IPI= IndTmed\cdot\left( IndBf+ IndPrp \right)$$

$$if Tmin>9^{\circ}C\underset{\to}{} Se IndTmed\cdot\left( IndBf+ IndPrp \right)<IndTmed\cdot IndUr\underset{\to}{}IPI= IndTmed\cdot IndUr$$

- **Magarey Model** (Magarey et al. 1991). To activate the transport of the inoculum from the soil to the plant, the model considers two conditions: 10 mm of rain and a temperature of at least 10°C for the last 24 hours. If these two simple conditions are met, the oospores laying in the soil must remain wet for at least 16 hours at temperatures equal or higher than 8°C. If also this occurs, the oospores germinate to macrosporangia and zoospores. At this point (first and second sets of conditions met), if the plant leaves remain humid for 45 degrees-hour, every precipitation event can favour the migration of macrosporangia and zoospores to the plant, eventually triggering infections.
- **UCSC Model** (Rossi et al. 2008). The model calculates at an hourly timesteps the main phases of the pathogen’s life cycle. Oospores created at the end of the vegetative season vernalize in the soil bed and in the first soil layers and during winter they reach morphological maturity. Germination of mature oospores can only happen when favourable environmental conditions trigger the dormancy-break and persist to allow germination. The newformed sporangia can survive for a variable time, e.g., from a couple of hours to multiple days, and release zoospores in the environment when water is available. Oospores survival is strongly linked to the presence of a water film and they can be dispersed in the environment (in the soil or on the plants) through rainfall. If the oospores reach a leaf, they ‘swim’ towards the stomata where they enter inside the leaf tissue. From here, hyphae colonize the host and after some time the first symptoms appear.

The model considers the following processes: the dormancy-break of the oospores, the length of the primary inoculation period, the density of oospores cohorts, oospores germination and sporangia appearance, sporangia survival, zoospores release and survival, zoospores infections, and the incubation period until the first symptoms become visible. To further delve into the model formalism, we refer to the original article (Rossi et al. 2008).

- **Laore Model**. It was developed by the rural development programs regional agency of Sarinia, Italy and is already implemented and used in the MISFITS-DSS platform.
- **Misfits Model** (Bregaglio et al. 2022). It is the model used in the MISFITS-DSS platform and predicts the life cycle of grapevine downy mildew.
