## Supplementary S3 for "Releasing the *octoPus,* an open-source digital tool to promote Integrated Pest Management"

Supplementary S3. Phenology calibration

Table S3.1 Model errors (RMSE, days) in simulating main phenological phases in the eight NUTS2 units used in calibration and evaluation.

| **Growth**  **stage** | **BBCH code** | **MAR** | **ABR** | **BAS** | **EMI** | **LIG** | **LOM** | **SAR** | **VEN** | **mean** |
| --- | --- | --- | --- | --- | --- | --- | --- | --- | --- | --- |
| Leaf development | 11 | 12.0 |  | 14.7 | 17.7 | 15.5 | 20.2 | 3.9 | 10.3 | 13.5 |
|  | 12 | 10.9 |  |  |  |  | 32.0 |  | 5.5 | 16.1 |
|  | 13 | 8.0 |  |  | 3.7 |  | 12.7 |  |  | 8.1 |
|  | 14 |  |  |  | 2.2 |  | 12.0 |  |  | 7.1 |
|  | 15 |  |  | 7.0 | 7.9 | 6.0 | 9.0 | 13.2 | 7.8 | 8.5 |
|  | 16 | 3.5 |  |  |  |  | 24.5 |  |  | 14.0 |
|  | 19 | 4.5 |  |  | 26.0 | 11.5 |  |  |  | 14.0 |
| Inflorescence emerge | 53 | 10.6 |  | 6.0 | 8.1 | 10.7 | 17.3 | 8.4 | 9.0 | 10.0 |
|  | 55 | 9.9 |  | 9.4 | 7.3 | 12.7 | 15.5 | 7.4 | 6.3 | 9.8 |
|  | 57 | 7.8 |  | 10.7 | 8.9 | 12.6 | 23.0 |  | 5.7 | 11.4 |
| Flowering | 60 | 6.0 |  | 28.0 | 7.5 |  | 18.5 |  | 17.0 | 15.4 |
|  | 61 | 9.4 | 11.4 | 12.8 | 5.5 | 11.8 | 22.0 | 7.4 | 9.3 | 11.2 |
|  | 63 |  |  | 2.0 | 3.2 | 10.3 | 3.0 |  |  | 4.6 |
|  | 65 | 10.9 |  | 12.0 | 8.0 | 5.0 | 12.2 | 1.5 |  | 8.3 |
| Development of fruits | 71 | 4.5 |  | 3.8 | 5.2 | 7.6 | 6.5 | 6.4 | 5.5 | 5.6 |
|  | 73 | 19.3 |  | 2.3 | 12.5 | 12.0 | 26.8 | 6.2 |  | 13.2 |
|  | 77 | 9.0 | 8.1 | 12.7 | 8.7 | 13.1 | 12.8 | 11.2 | 8.6 | 10.5 |
|  | 79 | 9.6 |  |  | 12.7 | 11.7 | 11.5 | 13.0 |  | 11.7 |
| Ripening of berries | 81 | 17.7 | 15.1 | 14.3 | 15.1 | 15.6 | 10.8 |  | 8.7 | 13.9 |
|  | 83 | 16.3 |  | 13.0 | 10.5 | 14.6 | 12.0 | 11.2 |  | 12.9 |
|  | 85 | 25.1 |  |  | 18.6 |  | 26.2 | 29.5 |  | 24.8 |
|  | mean | 10.8 | 11.5 | 10.6 | 10.0 | 11.4 | 16.4 | 9.9 | 8.5 |  |

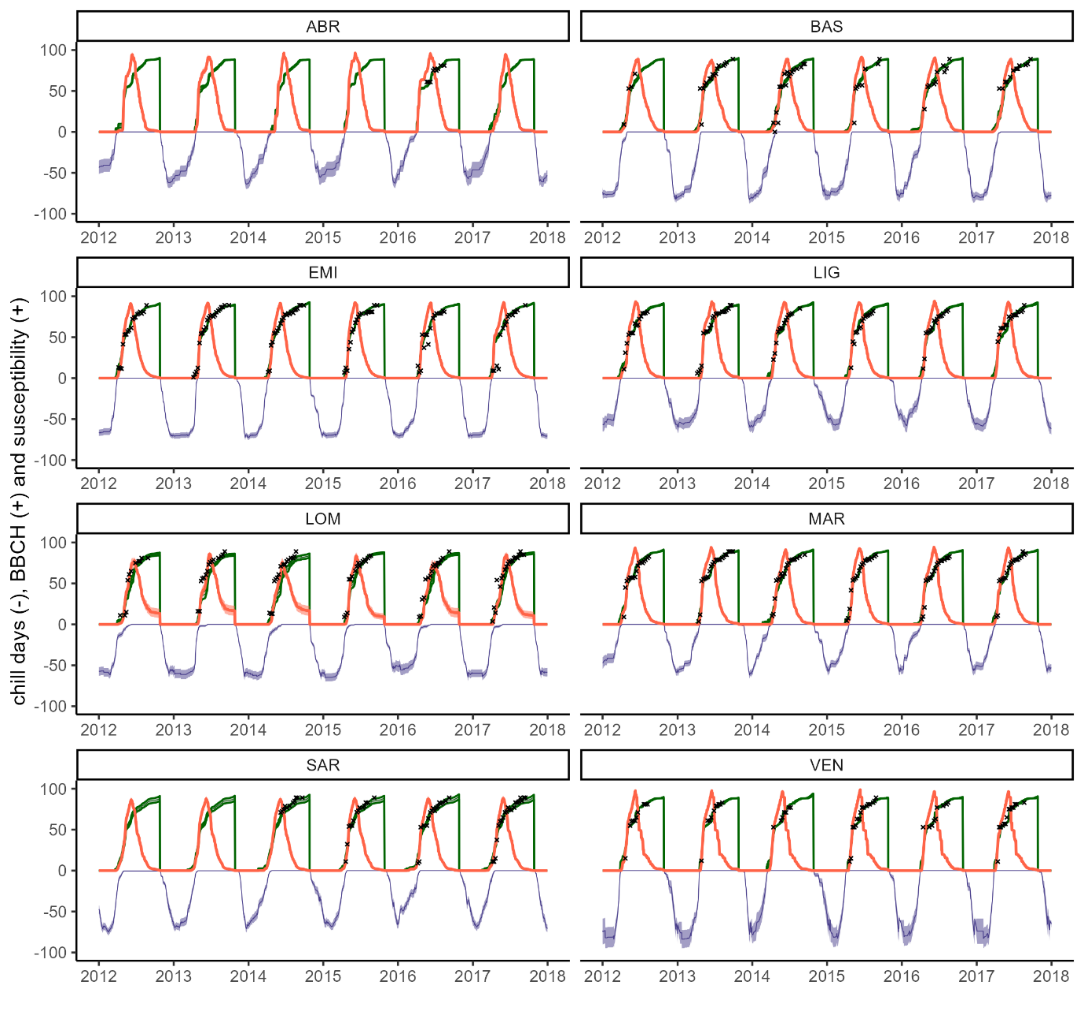

Figure S3.2 Phenology and host susceptibility dynamics simulated in the eight Italian regions where reference BBCH data were available. Chill and anti-chill days corresponding to the grapevine dormancy phase (blue, negative side of y-axis) and BBCH phases (green, positive side of y axis). The crosses indicate the reference BBCH observations. The orange lines correspond to host.
