## Supplementary S4 for "Releasing the *octoPus,* an open-source digital tool to promote Integrated Pest Management"

Supplementary S4. Results of the principal component analysis

Table S4.1. Principal components’ explained variance

| PC | eigenvalue | variance (%) | Cumulative |
| --- | --- | --- | --- |
| 1 | 3.05 | 50.9 | 50.87 |
| 2 | 1.87 | 31.2 | 82.1 |
| 3 | 0.54 | 8.9 | 91 |
| 4 | 0.33 | 5.4 | 96.4 |
| 5 | 0.19 | 3.1 | 99.6 |
| 6 | 0.03 | 0.4 | 100 |

Table S4.2. Coordinates of the variables on the first two Principal Components. (PC) Prec_tot = cumulated precipitation (mm), LW_tot = cumulated leaf wetness (hours), Tx_mean = mean maximum temperature (°C), Tn_mean = mean minimum temperature (°C), RHx_mean = mean maximum relative humidity (%), RHn_mean = mean minimum relative humidity (%). The variables have been computed from March to July.

| Variable | Unit | PC 1 | PC 2 |
| --- | --- | --- | --- |
| Prec_tot | mm | 0.66 | 0.60 |
| LW_tot | hours | 0.65 | 0.65 |
| Tx_mean | °C | -0.72 | 0.64 |
| Tn_mean | °C | -0.49 | 0.80 |
| RHx_mean | % | 0.86 | -0.15 |
| RHn_mean | % | 0.84 | 0.18 |

Table S4.3 Mean values of the variables used in the Principal Component Analysis within the four climate clusters from hierarchical clustering.

| Variable | Unit | Hot-Dry | Mild-Dry | Mild-Wet | Cold-Wet |
| --- | --- | --- | --- | --- | --- |
| Tx_mean | °C | 24.2^***^ | 22.5^**^ | 22.1^***^ | 17.6^***^ |
| Tn_mean | °C | 14.7^***^ | 13.5^*^ | 13.1 | 8.0^**^ |
| RHx_mean | % | 84.4^***^ | 87.4^*^ | 87.8^***^ | 90.8^***^ |
| RHn_mean | % | 45.6^***^ | 50.9 | 51.2^***^ | 51.7 |
| LW_tot | hours | 511.2^***^ | 752.6 | 986.6^***^ | 470.2^***^ |
| Prec_tot | mm | 158.1^***^ | 269.1 | 385.3^***^ | 171.8^***^ |
