## Supplementary S5 for "Releasing the *octoPus,* an open-source digital tool to promote Integrated Pest Management"

Supplementary S5. Performances of the Random Forest model

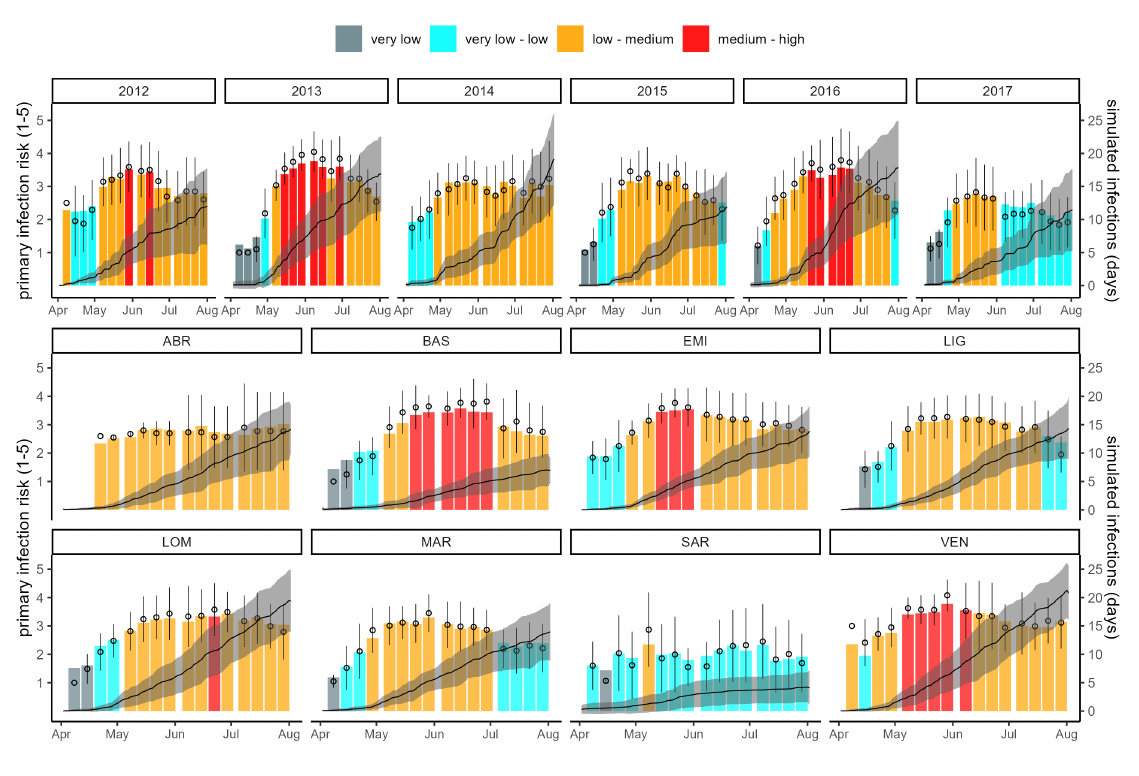

Figure S5.1 Performance of the Random Forest model (colored histogram) compared with reference infection risks (points and error bars) grouped by years (2012-2017, in the top row) and eight NUTS2 regions of italy, i.e., Abruzzo, Basilicata, Emilia-Romagna, Liguria, Lombardia, Marche, Sardegna, and Veneto. The black line is the median of the infections simulated by the eigth models and the ribbon corresponds to ± one standard error.
